# Comparison of DNA metabarcoding and morphological identification for stream macroinvertebrate biodiversity assessment and monitoring

**DOI:** 10.1101/436162

**Authors:** Joeselle M. Serrana, Yo Miyake, Maribet Gamboa, Kozo Watanabe

**Author notes:** corresponding author, Department of Civil and Environmental Engineering, Ehime University, Bunkyo-cho 3, Matsuyama, 790-8577, Japan.

## Abstract

Conventional morphology-based identification is commonly used for routine assessment of freshwater ecosystems. However, cost and time efficient techniques such as high-throughput sequencing (HTS) based approaches may resolve the constraints encountered in conducting morphology-based surveys. Here, we characterized stream macroinvertebrate species diversity and community composition via metabarcoding and morphological analysis from environmental samples collected from the Shigenobu River Basin in Ehime Prefecture, Japan. We compared diversity metrics and assessed both approaches’ ability to evaluate the relationship between macroinvertebrate community and environmental variables. In total, we morphologically identified 45 taxa (3 families, six subfamilies, 31 genera, and five species) from 8,276 collected individuals from ten study sites. We detected 44 species by metabarcoding, with 35 species collapsed into 11 groups matching the morphologically identified taxa. A significant positive correlation between logged depth (number of HTS reads) and abundance of morphological taxa was observed, which implied that quantitative data can be used for subsequent analyses. Relatively higher estimates of alpha diversity were calculated from the metabarcoding data in comparison to morphology-based data. However, beta diversity estimates between metabarcoding and morphology data based on both incidence and abundance-based matrices were correlated proving that community differences between sampling sites were preserved in the molecular data. Also, both models were significant, but metabarcoding data (93%) explained a relatively higher percentage of variation in the relationship between community composition and the environmental variables than morphological data (91%). Overall, we present both the feasibility and limitations of HTS-driven estimations of taxonomic richness, community composition, and diversity metrics, and that metabarcoding was proven comparable and more sensitive against morphology-based analysis for stream macroinvertebrate biodiversity assessment and environmental monitoring.

## 1. Introduction

Reliable and comprehensive, but rapid and cost-effective methods for monitoring freshwater ecosystems are encouraged due to the increasing threats of biological degradation faced by freshwaters worldwide (Carrizo et al., 2017). Since many ecological processes, stream characteristics and nutrient concentrations are important determinants of macroinvertebrate community composition (Heino, 2014; Shearer et al., 2015), macroinvertebrates have been the most commonly used focal groups for biological monitoring of the environmental quality of freshwater ecosystems (Menezes et al., 2010). They serve as good indicators of ecosystem health due to their high diversity, and different sensitivity to a range of natural and anthropogenic disturbances, which has been used to develop biotic indices for extensive monitoring programs (Aylagas et al., 2016).

Conventional morphological analysis is most commonly used in routine monitoring programs evaluating environmental quality changes. However, this is not only time consuming but has serious issues with accuracy, and consistency in the level of taxonomic identification that highly depends on taxonomic expertise (Hajibabaei et al., 2011). Specifically, small organisms such as the larval stages of stream macroinvertebrates frequently used for river biomonitoring are often difficult or impossible to identify at finer taxonomic resolution (e.g., species level) (Sweeny et al., 2011). A promising alternative approach is DNA metabarcoding - a combination of amplicon-based high-throughput sequencing (HTS) analysis and DNA taxonomy (Hebert et al., 2003). High-throughput amplicon sequencing can process large number of individuals simultaneously and in parallel (Thudi et al., 2012) making it faster and cheaper than the conventional Sanger sequencing (Voelkerding et al., 2009). Following a comprehensive read processing step, most metabarcoding pipelines carry out taxonomic assignments by comparing clustered reads or operational taxonomic units (OTUs) to a reference sequence database such as Genbank (Benson et al., 2012) and the Barcode of Life Data System (BOLD) (Ratnasingham and Hebert, 2007). Metabarcoding promises cost-effective and quicker assessments with more comprehensive and verifiable taxonomic identification that is less reliant on taxonomic expertise (Baird and Hajibabaei, 2012; Yu et al., 2012; Emilson et al., 2017).

The application of HTS-based approaches for biodiversity assessments has been rapidly expanding across a wide range of fields, including the biomonitoring of stream macroinvertebrates (Baird and Hajibabaei, 2012; Beng et al., 2016). Previous studies have assessed the ability of DNA metabarcoding to identify macroinvertebrate communities in parallel to morphology-based identification. DNA metabarcoding provides broader taxonomic coverage and finer resolving power (Soininen et al., 2015). Hence, species that may exhibit diverse environmental responses would have a higher chance of detection benefiting study systems that require species-level identification (Elbrecht et al, 2017; Carew et al., 2018). With this advantage, DNA metabarcoding may provide stronger discriminatory power in detecting environmental variables that influence community composition compared to traditional methods (Emilson et al., 2017).

Then again, amplicon-based HTS analysis for biomonitoring has technical difficulties associated with the quantitative assessment of the abundance of each species in a community (Elbrecht and Leese, 2015). Quantification of relative abundance is useful for community characterization, and assessment of biological indices as most diversity measures depend on a reliable recovery of taxonomic abundances (Gotelli and Chao, 2013; Aylagas et al., 2014; Bucklin et al., 2016). In theory, highly abundant species will yield higher concentrations of template DNA in the community DNA soup (Yu et al., 2012) leading to a positive correlation between the number of HTS reads (depth) and the abundance of species. However, interspecific differences in PCR primer compatibility and body mass may affect the efficiency of PCR amplification, potentially collapsing this correlation (Kowalczyk et al., 2011, Deagle et al., 2009, Zhou et al., 2013). Most metabarcoding studies were based on small organisms, such as algal, bacterial, and planktonic communities, where the morphological quantification of abundance is difficult or impossible, and so abundance and depth cannot be directly compared. However, recent metabarcoding studies have tested and found such positive correlation using relative abundances of morphologically identified taxa. Most of these studies were taxonomically limited to the family Nematoda (Porazinska et al., 2010), Chironomidae (Carew et al., 2013) or calanoid copepods (Clarke et al., 2017) that include similarly-sized species, while a handful tested environmental samples composed of macroinvertebrate taxa with varying size (e.g., Aylagas et al., 2016; Elbrecht et al., 2017; Krehenwinkel et al., 2017; Serrana et al., 2018). However, more tests with wider taxonomic and body size ranges such as macroinvertebrate communities are necessary to verify this relationship for taxonomically diverse communities for use in stream assessment.

In this study, we compared conventional morphological identification against DNA metabarcoding to explore the feasibility and limitations of HTS analysis for stream macroinvertebrate diversity assessment and biomonitoring. Environmental samples of stream macroinvertebrate communities were collected from the Shigenobu River in Shikoku Island, Japan. First, we assessed the relationship between the number of HTS-reads (depth) and relative abundance of taxonomic groups from the metabarcoding and morphology-based data to verify the applicability of abundance-based metrics for subsequent analyses. We then compare alpha and beta diversity metrics calculated from the two data sets and examine how HTS-derived data compares with the traditional method for assessment. Finally, we evaluated the ability of metabarcoding and morphological surveys in assessing the influence of stream environmental conditions in macroinvertebrate community composition in order to assess the capability of metabarcoding data for stream environmental monitoring.

## 2. Materials and Methods

### 2.1. Study area and sample collection

Field survey was conducted in the Shigenobu River basin in Ehime Prefecture, Japan (33°47′N, 132°47′E; Fig. 1) in August 2012. The basin has an area of approximately 445 km2, and the river originates from 1,232 m above sea level in its headwaters, flows 36 km along the length of the corridor, and finally discharges into the Seto Inland Sea as a fifth order river. The mountainous area is covered by plantation and secondary forests, while the lowland area predominantly consists of urban and agricultural land. Annual precipitation is approximately 1,300 mm with the wet season occurring in summer. From the mid-to lower reaches of the river, the channel is braided (bankfull width approximately 300 m maximum), and the flow tends to be intermittent owing to a thick channel layer of alluvial deposits. At present, there are three intermittent reaches in the river (upper: 18–22 km from the mouth; middle: 10–16 km; and lower: 4.9–7.4 km) (Kawanishi et al., 2011). Macroinvertebrate samples were collected from 10 sites longitudinally located from the headwater to the lower reaches of the Shigenobu River (elevation range: 4–405 m; Table 1, and Fig. 1). We did not conduct sampling within the upper and middle intermittent reaches owing to the loss of surface water. Three equally spaced transects (40-m intervals) were established in each study site, and the macroinvertebrates were sampled at the center of each transect using a Surber sampler (25 × 25 cm quadrat, 0.5-mm mesh). The samples were immediately preserved in 99.5% ethanol, which was replaced twice in the field to prevent DNA degradation. The collected macroinvertebrates were then sorted and morphologically identified to the lowest taxonomic level possible using the taxonomic keys of Kawai and Tanida (2005).

**Figure 1.**
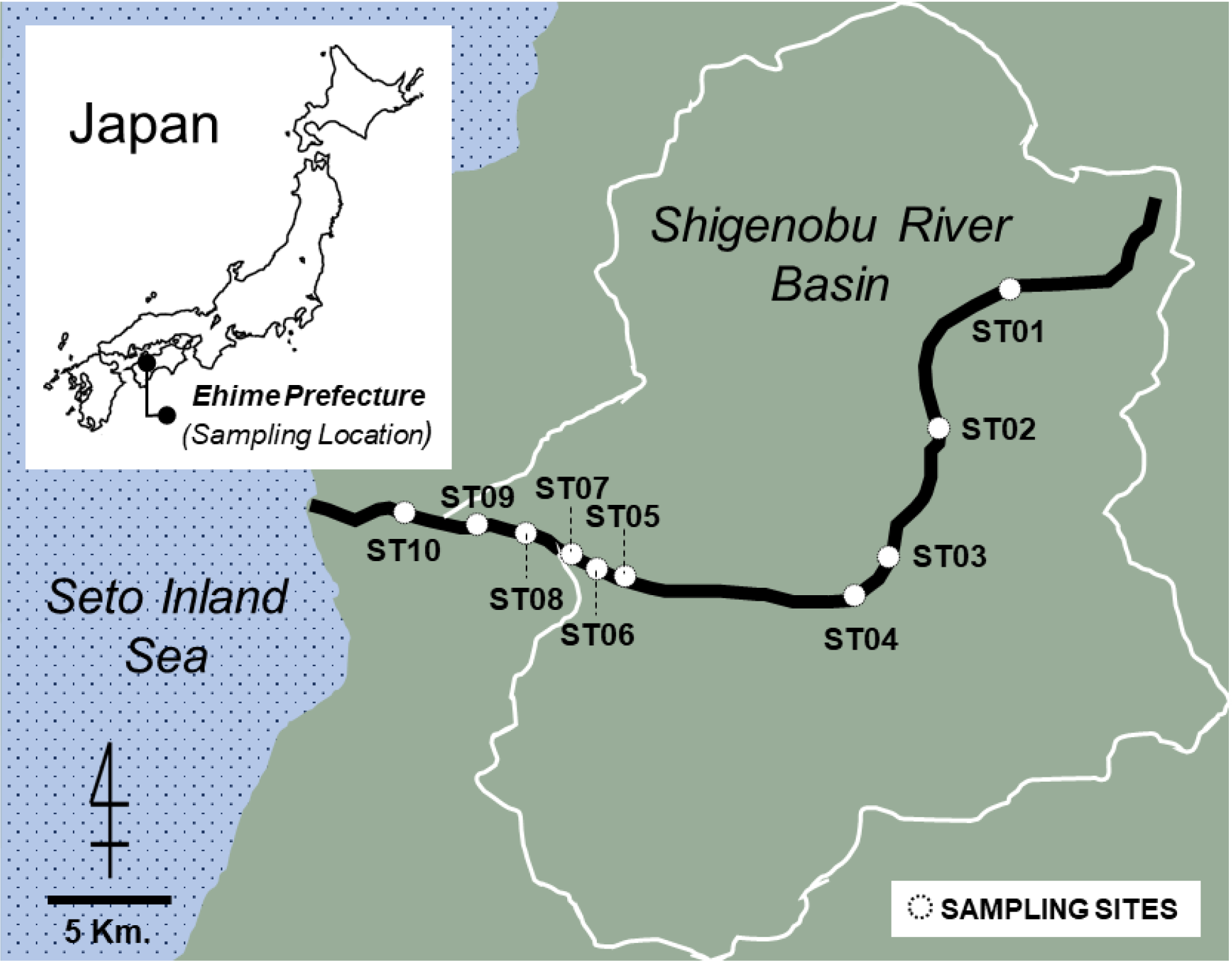
Map of the ten study sites along the Shigenobu River in Shikoku Island, Japan. The altitude of each site is provided in Table 1.

### 2.2. Library preparation and 454 pyrosequencing

All specimens collected per study site were pooled, and dry weight (DW) was measured using a UMX2 Ultra-microbalance (Mettler-Toledo, Inc., USA). Based on the measured DW (19 – 403 mg per site), the samples were separated into ≤ 10 mg DW portions and placed in 1.5-ml tubes. The samples were then homogenized in the tubes using pestles. DNA was extracted using DNeasy® Blood and Tissue Kit (Qiagen GmbH, Hilden, Germany) following the manufacturer’s instructions. Extracted DNA quantity and quality were measured using a NanoDrop 2000 spectrophotometer (Thermo Scientific, Wilmington, DE, USA). DNA originating from the same study site were mixed with equal amounts of volume. A 658-bp fragment of the Cytochrome Oxidase I (COI) DNA barcode region was amplified using the universal primers LCO-1490 and HCO-2198 (Folmer et al., 1994), which both had 454-specific fusion primers (11-mer) and a 4-mer key sequence (TCAG) added to the 5′-end. To pool multiple PCR products from the 10 study sites in one 454 pyrosequence run, a Molecular Identifier (MID) tag (6-mer) was also added to the 5′-end of the LCO-1490 primer. PCR was conducted with a 40-μl reaction volume containing 20 μl of Phusion® High-Fidelity PCR Master Mix with hydrofluoric acid (HF) buffer (New England Biolabs, UK), 2 μl of each PCR primer (10 uM), 3 μl of template DNA, and 13μl of PCR-grade water using a T100TM Thermal Cycler (Bio-Rad Laboratories, CA, USA). The PCR conditions were denaturation at 94°C for 2 min; 30 amplification cycles at 94°C for 30 s, 40°C for 30 s, and 72°C for 60 s; and final extension at 72°C for 10 min. Two (duplicate) PCR products were generated for each site. The resulting 20 PCR products were visualized using gel electrophoresis, and the target bands of successful samples were sliced and purified using the QIAquick® Gel Extraction Kit (QIAGEN, Germany). The DNA concentration of each purified PCR product was measured with a Quantus™ Fluorometer (Promega, Wi., USA) using the QuantiFluor® dsDNA System (Promega, Wi., USA) and was mixed with an equal molar amount. Pyrosequencing was carried out by Hokkaido System Science Co., Ltd. (Sapporo, Japan) using the 1/16 region of a GS FLX Titanium instrument (454; Roche Diagnostics).

### 2.3. Bioinformatics analysis

In total, 165,508 passing filter reads with an average length of 328-bp were acquired. FastQC v0.11.5 (Andrews 2010) was used to assess sequence quality. The raw pyrosequencing data were initially processed using Trimmomatic v0.36 (Bolger et al., 2014) to remove non-biological sequences, i.e., primer and index sequences. Quality filtering of the reads was performed following the UPARSE pipeline (Edgar 2013) using the maximum expected error parameter. Reads with >1 maximum expected error and length <200-bp were discarded. Surviving reads for the ten sites were truncated to 200-bp to obtain globally aligned reads, pooled and clustered into operational taxonomic units (OTU) using USEARCH v9.2.64 (Edgar 2010). For *de novo* OTU assembly, the reads were dereplicated into unique sequences with 100% similarity. Resulting unique sequences were then clustered into operational taxonomic units (OTUs) with a similarity cut-off value of 97%, discarding putatively chimeric and singleton sequences. BLAST searches were performed on the OTU representative sequence against reference databases, i.e., Barcode of Life Database (BOLD) and GenBank. Taxonomic identification was assigned based on the best BLAST hit to a sequence with ≥97% identity, e-value ≥10^−5^ and minimum query coverage >90%. Representative OTU sequences without significant BLAST hits and non-arthropod matches were excluded from subsequent analyses.

To examine the correlation between sequence depth and abundance, we plotted the number of reads of given taxa identified by BLAST against the abundance (individuals per 0.19 m^2^) of the morphologically identified taxa (morpho-taxa). Metabarcoding identified the samples at the species level while the morpho-taxa were identified at inconsistent taxonomic levels (e.g., *Hydropsyche orientaris* vs. *Hydropsyche*). The identified metabarcoding species were collapsed into a coarser taxonomic level (e.g., *Hydropsyche*) or “meta-taxa” to facilitate a balanced comparison between the two data sets. Detected metabarcoding taxa that did not match the morpho-taxa or false positive detection was retained at the species level in the dataset.

### 2.4. Biodiversity analysis

Diversity was evaluated within-community (alpha diversity) and between the communities (beta diversity) for both morpho-taxa and metabarcoding-identified species. Quantitative Insights into Microbial Ecology (QIIME) (Caporaso et al., 2010) was used to estimate diversity, with data matrices rarefied to the sampling site with the lowest abundance to equalize the number of reads or individuals. Alpha metrics assessed were Chao 1 richness, Fisher’s alpha, Simpson’s index, and the Shannon-Wiener diversity index. Linear regression analyses were performed on log-transformed values to test the correlation of each alpha diversity matrices between the two data sets. Non-phylogenetic beta diversity was estimated using both qualitative metric (measure changes in communities based on presence or absence/incidence) – Binary-Jaccard dissimilarity, and quantitative metric (measure differences in relative abundances between communities) – Bray-Curtis dissimilarity. Mantel test was used to test the correlation between the morpho-taxa, and metabarcoding-identified species data, and Principal Coordinates Analysis (PCoA) to visualize the dissimilarity matrices. Procrustes test was used to test for correlation between the two ordinations.

### 2.5. Relationship between environmental variables and macroinvertebrate composition

Physical and chemical characteristics were collected from each study sites. Physical habitat characteristics were measured at three equally placed points along five transects of each study sites. Stream characteristics such as width were measured on each transect (n = 5), while stream depth was measured to the nearest 1 cm at each point of the transects (n = 15). Current velocity was measured above the streambed at each point using a portable current meter (Model CR-7WP; Cosmo-Riken Inc., Osaka, Japan). Electric conductivity and dissolved oxygen were measured using a multiparameter water quality meter (Model 556MPS, YSI Inc. Yellow Springs, OH, USA). Substrate coarseness and embeddedness were evaluated by visual assessment (Matthaei et al., 1999; Miyake and Akiyama, 2012). Single surface water samples were collected midstream of each site for water chemistry analysis. Periphyton biomass was estimated by measuring chlorophyll *a* concentration. The quantity of coarse particulate organic matter (CPOM) contained in each sample was estimated via ash-free dry mass (AFDM, g m^−2^). Total nitrogen (TN) and total phosphorus (TP) were measured following standard methods (Apha, 2005). Redundancy analysis (RDA) was performed and plotted in the *vegan* R package (Oksanen et al., 2014) to visualize the relationships between the physical-chemical characteristics and the macroinvertebrate taxa detected via morphological and metabarcoding identification. Variance inflation factors (VIF) of the environmental variables were checked using the *vif* function in the *faraway* R package (Faraway, 2016). Variables selected have VIF values <4 and tolerance >0.20 variables to avoid issues of multicollinearity. ANOVA was run with 10000 permutations to assess the significance of constraints. Analyses were performed for the model (global test), for each constrained axis (setting: by = “axis”), and for each predicting variables (setting: by = “margins”) of the two data sets. Morphological and metabarcoding data were Hellinger-transformed before ordination. The predicting variables were log-transformed to meet the assumptions of normality and equal variance. Statistical analyses were run in R v.3.3.1.

## 3. Results

### 3.1. Taxonomic identification

A total of 8,276 individuals and 45 morpho-taxa (3 families, 6 subfamilies, 31 genera, and 5 species) were collected from the ten study sites (Table 1). The top three most dominant morpho-taxa were Chironominae (3,144 individuals, 38%), *Baetis* (2,203 individuals, 26.6%), and Orthocladiinae (1,639 individuals, 19.8%). The 454 pyrosequencing analysis generated a total of 165,508 reads (range: 8,593 – 31,291 reads/site; mean: 19,942 reads/site) with an average length of 328-bp (range: 31 - 594-bp). Raw sequence data is available from NCBI Sequence Read Archive (SRA) with an accession number SRR7957429. After quality filtering, 81,836 reads (49.5%) were retained, with 79,902 reads (48.3%) mapped to 156 OTUs. No significant matches in the BOLD database were obtained. However, 53 OTUs (34%) consisted of 47,641 reads (59.6% of reads mapped to OTUs) had significant BLAST hits to arthropod sequences in GenBank with ≥97% identity and e-value ≥10^−5^ identified to the species level. The remaining OTUs either have BLAST identity under the matching criteria (91 OTUs, 31,962 reads), non-arthropod sequence match (7 OTUs, 171 reads), or no match (5 OTUs, 128 reads). Metabarcoding identified 44 species (4 matches with “*sp*.”) under 6 orders, 13 families, and 29 genera. For the orders, Diptera was the most abundant with 30,955 reads (65% of the taxonomically-identified arthropod sequences), followed by Ephemeroptera and Trichoptera with 14,881 reads (31.2%) and 1,570 reads (3.3%) respectively. Other orders, i.e., Plecoptera, Odonata, and Podocopida, had <1% read abundance. See Fig. S2 for the relative abundance of the metabarcoding-detected species, and the meta- and morpho-taxa.

Significant positive correlations were found between the total abundance of morpho-taxa and the read abundance of meta-taxa for all sites, both in analyses including (R^2^ = 0.18; *p* = 0.001) and excluding (R^2^ = 0.48; *p* = 0.02) false positive and false negative detections (Fig. 2). Positive linear correlations (*p* < 0.05) were also found for each sampling site including false positive and false negative detections, except for four sites i.e. sites 3, 4, 5 and 7 (Table 1). Thirty-five species (95.5% of reads) from our metabarcoding data matches or were under the taxonomic identification of 11 morpho-taxa (92.9% of individuals). The remaining 9 species (4.5% of reads) were false positive detections, while 34 morpho-taxa (7.12% of individuals) were undetected.

**Figure 2.**
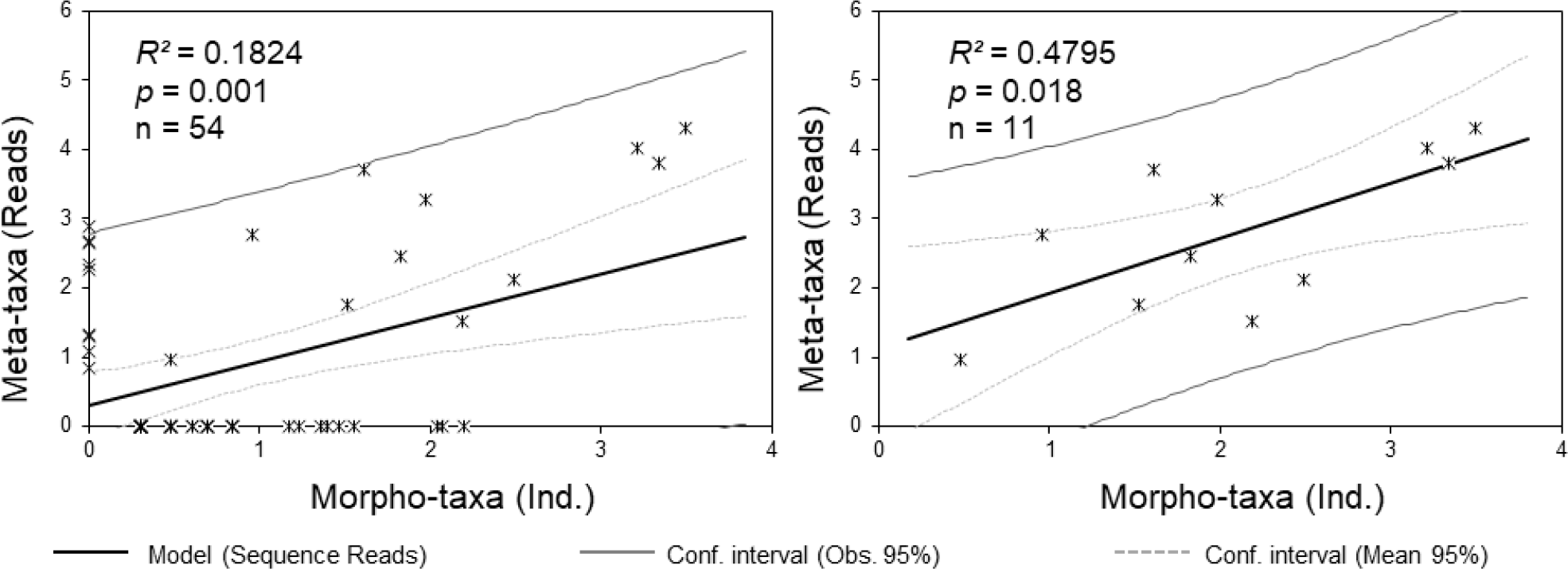
Correlation between the log sample abundance (morphologically-identified taxa/morpho-taxa) and log 454-read abundance (metabarcoding-identified taxa) of all sites showing analysis including (left) and excluding (right) false positive and false negative detection.

**Table 1.** Summary of altitude, DNA metabarcoding data, and species richness across the ten study sites.

| Study site | Sample Size | Read Processing |  |  | Metabarcoding |  |  | Morpho-taxa | Meta-taxa (Match) | False Positive | False Negative | Correlation Analysis |  |  |
| --- | --- | --- | --- | --- | --- | --- | --- | --- | --- | --- | --- | --- | --- | --- |
|  |  | Raw Reads | Reads Mapped to OTUs | OTUs | Arthropod Reads<sup>a</sup> | Meta-species | Meta-taxa |  |  |  |  | Pr > F<sup>b</sup> | R<sup>2</sup> | R |
| Site 01 | 603 | 18,867 | 10,593 | 14 | 4,338 | 13 | 3 | 5 | 2 | 1 | 3 | 0.03 | 0.73 | 0.85 |
| Site 02 | 228 | 31,300 | 15,371 | 15 | 11,704 | 14 | 6 | 7 | 3 | 3 | 4 | 0.03 | 0.48 | 0.69 |
| Site 03 | 208 | 8,593 | 4,941 | 24 | 2,520 | 20 | 8 | 9 | 5 | 3 | 4 | 0.10<sup>c</sup> | 0.24 | 0.49 |
| Site 04 | 551 | 15,033 | 4,797 | 14 | 3,987 | 12 | 9 | 12 | 5 | 4 | 7 | 0.49<sup>c</sup> | 0.03 | 0.18 |
| Site 05 | 1,347 | 12,598 | 5,792 | 28 | 3,534 | 25 | 9 | 14 | 6 | 3 | 8 | 0.08<sup>c</sup> | 0.19 | 0.44 |
| Site 06 | 1,051 | 31,291 | 14,848 | 30 | 11,837 | 24 | 10 | 16 | 6 | 4 | 10 | 0.01 | 0.34 | 0.58 |
| Site 07 | 539 | 17,510 | 7,304 | 24 | 4,957 | 20 | 12 | 20 | 8 | 4 | 12 | 0.21<sup>c</sup> | 0.06 | 0.25 |
| Site 08 | 2,740 | 10,626 | 4,944 | 26 | 2,802 | 21 | 12 | 25 | 8 | 4 | 17 | 0.01 | 0.25 | 0.5 |
| Site 09 | 843 | 8,784 | 5,106 | 28 | 1,462 | 22 | 9 | 22 | 6 | 3 | 16 | 0 | 0.39 | 0.62 |
| Site 10 | 166 | 10,906 | 6,206 | 15 | 500 | 13 | 6 | 27 | 5 | 1 | 22 | 0.03 | 0.17 | 0.41 |
| **Total** | **8,276** | **165,508** | **79,902** | **53** | **47,641** | **44** | **20** | **45** | **11** | **9** | **34** | **0.001** | **0.18** | **0.43** |

<sup>a</sup> arthropod sequence match at >97% identity and e-value >10<sup>-5</sup> identified to the species level in GenBank.
<sup>b</sup> including false positive and false negative detection.
<sup>c</sup> no significant correlation with p-value > 0.05.

Most of the undetected morpho-taxa (false negative) have low representation (<6 individuals) in the community sample, with only nine groups having >14 individuals (Table 2).

**Table 2.**
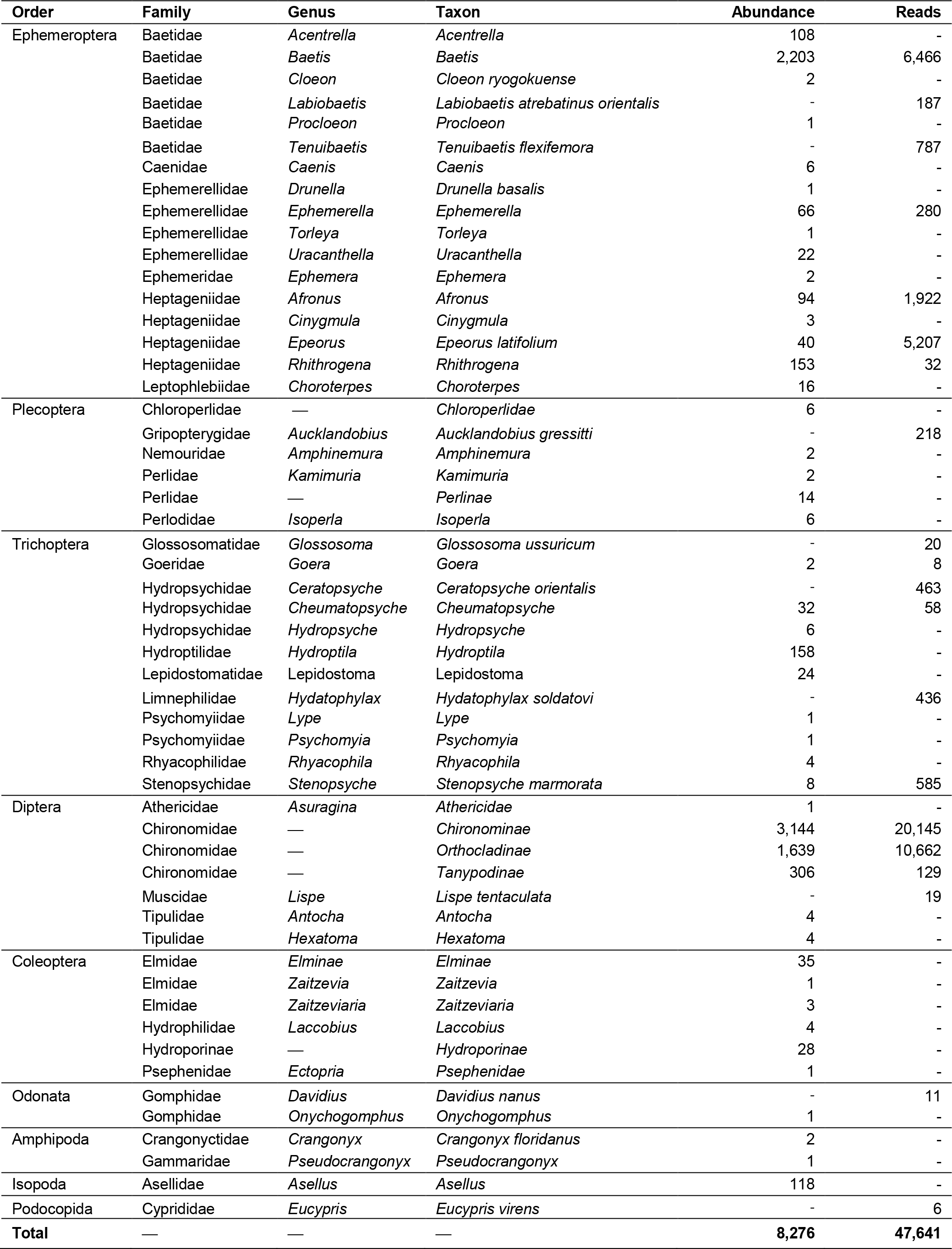
Absolute abundances of macroinvertebrates based on morphological identification (morpho-taxa) and metabarcoding sequence reads (meta-taxa). The correlations between these are shown in Figure 2.

### 3.2. Comparison between morphological and metabarcoding-based diversity metrics

Taxonomic richness or alpha diversity was assessed for both the morpho-taxa and metabarcoding-identified species data sets. Metabarcoding species data showed relatively higher values for Chao 1 richness and Fisher’s alpha, except for site 10, and sites 4 and 10, respectively. Additionally, Simpson’s and Shannon-Wiener indices have relatively higher values for some sites in comparison to the morpho-taxa dataset (Table S1). However, linear regression analysis revealed that the alpha diversity metrics of the two data sets were not significantly correlated i.e., Chao 1 (*p*-value = 0.458), Fisher’s alpha (*p*-value = 0.698), Simpson’s index (*p*-value = 0.506) and Shannon-Wiener index (*p*-value = 0.653). On the contrary, mantel testing both beta diversity distances, Binary-Jaccard (Mantel *r* = 0.4701, 9999 permutations, *p* = 0.0073) and Bray-Curtis (Mantel *r* = 0.4581, 9999 permutations, *p* = 0.0075) dissimilarities of the morpho-taxa and metabarcoding-identified species data matrices showed significant correlation. PCoA ordination plots of the beta diversity estimates were also significantly correlated shown via Procrustes analysis (Fig. 3).

**Figure 3.**
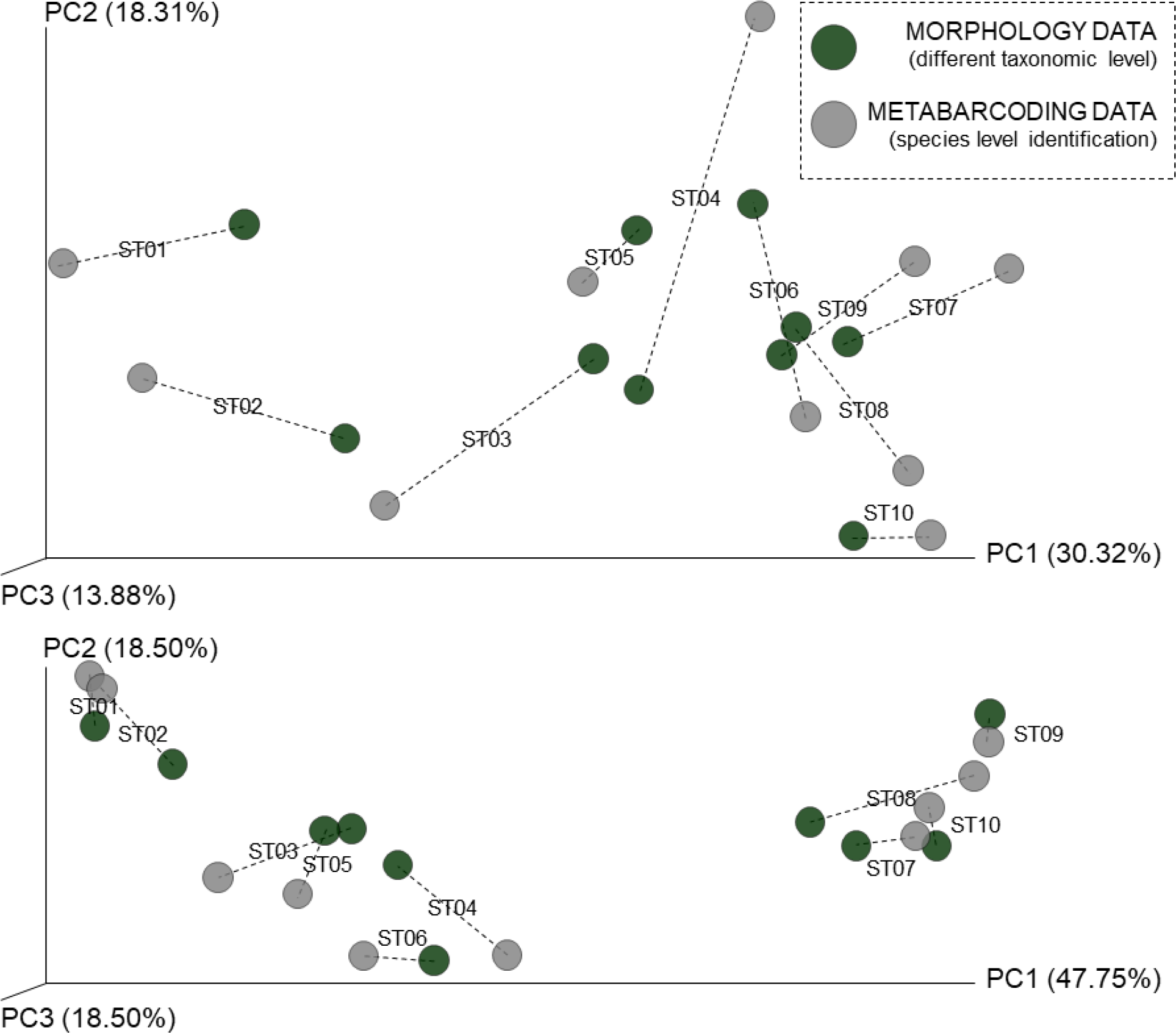
Principal coordinate analysis (PCoA) ordinations of beta diversity estimates (coords scaled by percent explained): Binary-Jaccard (Procrustes: *p*-value = 0.038) (top) and Bray-Curtis dissimilarity (Procrustes: *p*-value < 0.001) (bottom).

### 3.3. Environmental variables and macroinvertebrate community composition

The detailed physical-chemical characteristics collected across the ten study sites are presented in Table S1. Redundancy analysis (RDA) was performed to visualize the relationships between stream physical-chemical characteristics and macroinvertebrate community composition. Seven environmental variables were selected as predicting variables, i.e., chlorophyll *a*, conductivity, CPOM, discharge, dissolved oxygen, TN, and TP. Global models of the morphology (*p* = 0.035) and metabarcoding (p < 0.001) data set constrained by the environmental variables were found to be statistically significant following a permutation ANOVA test. RDA explained about 91% of the total variability between macroinvertebrate composition and the environmental variables for the morphology data. Of the 91%, 59% was explained by RDA1 and 15% by RDA2. However, only the first axis was statistically significant (RDA1: *p* = 0.033). For the metabarcoding-identified species data, RDA explained 93% of the total variability. From this, the first two statistically significant axes explained 44% and 19 % respectively (RDA1: *p* < 0.001; RDA2: *p* = 0.011). The first axes (RDA1) of both data sets were positively loaded with chlorophyll *a*, conductivity, TN, and TP (biplot scores = 0.5171, 0.5559, 0.8195, and 0.7151 respectively), and negatively loaded with CPOM, dissolved oxygen, and discharge (biplot scores = −0.5367, −0.7587, and −0.1013 respectively). For second axis (RDA2) of the metabarcoding data was positively loaded with CPOM, discharge, and TN (biplot scores = 0.0140, 0.2053, and 0.0.0651 respectively), and negatively loaded with chlorophyll *a*, dissolved oxygen, conductivity and TP (biplot scores = −0.3012, −0.6468, −0.2314 and −0.1642 respectively) (Fig. 4). Also, permutation ANOVA set by margin to assess the marginal effects of each marginal term analyzed in the model with all other variables showed all environmental characteristics, except for chlorophyll *a*, were statistically significant for the metabarcoding data (Table S3).

**Figure 4.**
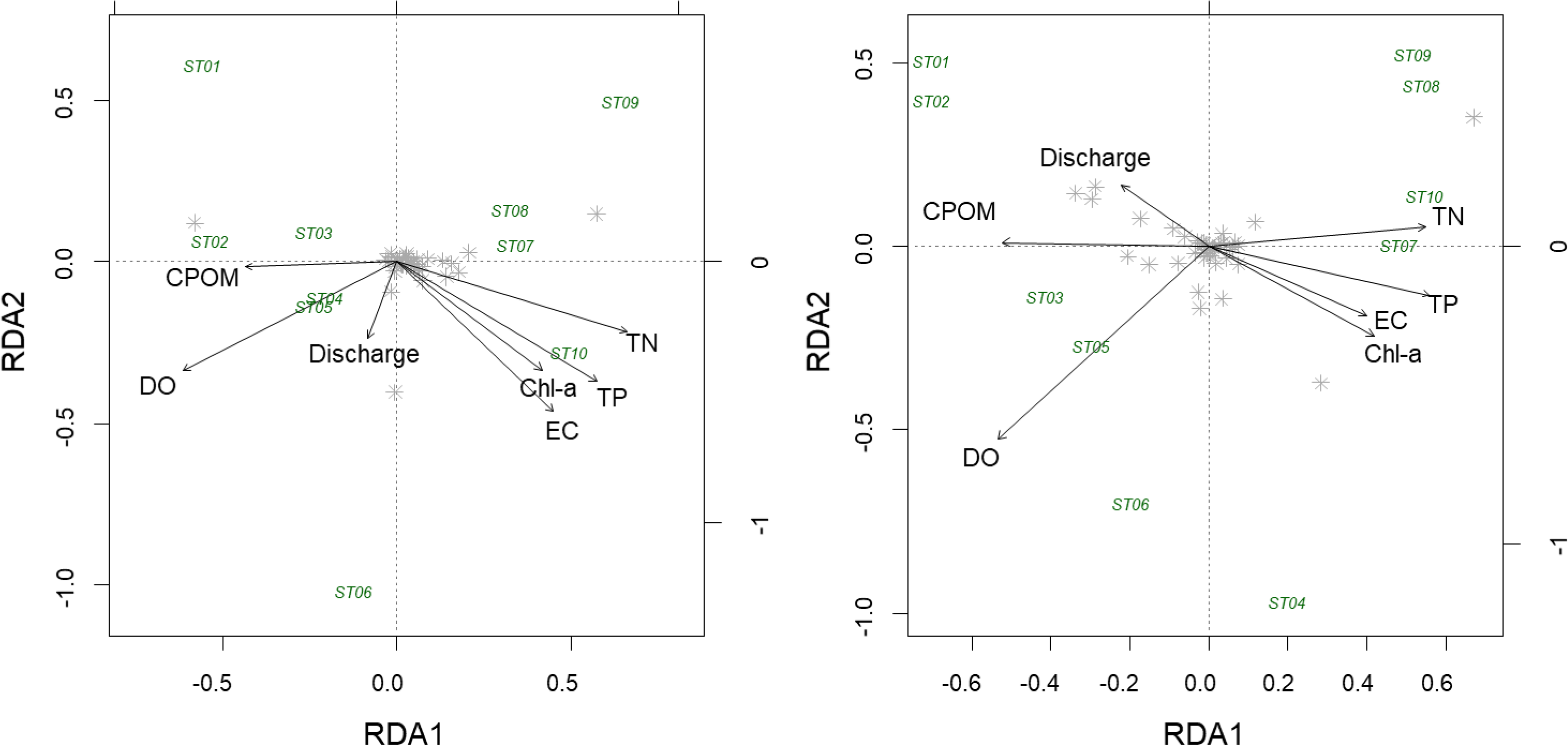
Redundancy analysis (RDA) ordination plot of the morphologically-identified taxa (left) and metabarcoding-identified macroinvertebrate species (right) constrained by environmental variables. Both global models were found to be statistically significant following a permutation ANOVA test (p = 0.035, p < 0.001 respectively). “Chl-a” stands for chlorophyll a, “CPOM” for the coarse particulate organic matter, “DO” for dissolved oxygen, “EC” for conductivity, “TN” for total nitrogen and “TP” for total phosphorus.

## 4. Discussion

Assessing the changes in community composition in response to environmental variability is a key aspect of biodiversity assessment and monitoring (Ives and Carpenter, 2007). We evaluated the ability of the morphological and metabarcoding approaches to assess the relationship between physical-chemical characteristics and macroinvertebrate community composition. Community-environment relationships can be quantified by explained variance from a redundancy analysis (Lu et al., 2016). Both models were significant, but the metabarcoding data set explained a relatively higher percentage of variation between the environmental variables and community composition. Although both approaches presented almost similar responses for each predictor variables (except for CPOM, discharge, and TN), only metabarcoding data ordination explained statistically significant variability between the environmental variables and community composition for the first two axes, and for each marginal terms. Hence, we can interpret that statistically significant physical-chemical characteristic from these analyses, i.e., conductivity, CPOM, discharge, dissolved oxygen, TN and TP strongly influenced macroinvertebrate community composition along the Shigenobu River basin during the conduct of the study.

Evaluation of the sufficient taxonomic level for biodiversity assessments are vital for the establishment of a cost-effective monitoring program since identification at a higher taxonomic level (e.g. family or phylum) will reduce cost and processing time, requiring lesser taxonomic expertise (Thompson et al., 2003). Choosing the most appropriate taxonomic resolution highly depends on the biological group, the environment of the site and the objectives of the study (Machado et al., 2015). However, taxonomic identification with coarser resolution groups together biological species with different environmental preferences, masking the relationship between taxonomic composition and environmental variables (Martin et al., 2016). Nonetheless, DNA metabarcoding can provide a much more accurate taxonomic identification with finer resolution, so preliminary assessment of the sufficient taxonomic resolution appropriate for a specific study would not be required.

We observed no significant correlation between the alpha diversity measures of morphological and metabarcoding data sets. The lack of correlation could be due to the difference in taxonomic resolution, with the latter having relatively higher taxonomic richness per site. Previous studies using metabarcoding have reported a generally higher species richness compared to morphology-based surveys (Cowart et al., 2015). DNA metabarcoding uncovered specific taxonomic groups undetected (false positive) from morphological assessments which magnifies the variation in community composition (Emilson et al., 2017). Molecular-based surveys have been proven to provide a broader view of the taxa present in the community due to its ability to uncover small sized, even eggs and juveniles of larger individuals from the collected environmental sample (Cowart et al., 2015). This, in addition to the undetected (false negative) morpho-taxa most likely resulted in the lack of correlation between the alpha diversity values of our morphological and metabarcoding-based approaches. On the contrary, significant correlations were observed for both beta diversity estimates assessed based on presence-absence (incidence) and sample/read abundance. Suggesting that changes in community composition observed from the metabarcoding data were congruent with the morphology-based assessment leading to similar interpretation about the ecological state of the sampled river. Our results were congruent with the report of Gibson et al. (2015) that HTS analysis provides a complete site-region-level biodiversity estimation (i.e., alpha, beta and gamma diversity metrics). These support previous studies that proclaimed HTS-based analysis is a suitable method for freshwater macroinvertebrate diversity assessments as an alternative to morphology-based surveys (Hajibabaei et al., 2011; Carew et al., 2018; Serrana et al., 2018).

Whereas many previous amplicon-based HTS studies have focused on microbial or planktonic taxa (e.g., Kermarrec et al., 2013, González-Tortuero et al., 2015, Hajibabaei et al., 2012, Tang et al., 2012, Zimmermann et al., 2015), we assessed environmental samples of diverse size with representatives having relatively large body size (>0.6 mm) which allowed us to directly compare the HTS data from conventional morphological identification. The positive correlation between the abundance of morphological taxa and sequence read abundance of the metabarcoding taxa among the merged taxonomic groups suggests that metabarcoding could be used to quantify relative species abundance based on depth information. It is worth to note that in accordance to the recent report of Elbrecht et al. (2017), we observed this correlation with a taxonomically broader community (i.e. the entire stream macroinvertebrate community) in comparison to former studies that also reported a positive correlation of sample and read abundance such as Nematoda (Porazinska et al., 2010), Chironomidae (Carew et al., 2013), and calanoid copepod (Clarke et al., 2017) communities that assessed similarly-sized species. Although we did not measure the biomass (dry mass or wet mass) of individual samples, we can assume that there was large body mass variation between the small, e.g., Chironomidae: reported mean individual dry mass = 0.06 mg (Watanabe et al., 2008) and large e.g. *Stenopsyche marmorata*: 14.26 mg (Nakagawa and Takemon, 2015) taxa in our samples.

Large taxa are expected to yield a larger portion of template DNA in the community DNA soup, leading to a higher number of reads regardless of their abundances. However, the fact that we were still able to detect an abundance-depth correlation demonstrates the robustness of this relationship. It is possible that the presumed high yield of template DNA from abundant species may overcome the potential bias effect of body mass variation among taxa. However, a comparison of the correlations between both body mass and abundance with depth is required in the future to validate these findings, considering the potential influence of taxon biomass to sequence reads (Elbrecht, Peinert, and Leese, 2017). This positive abundance-depth correlation also indicates that HTS may have low statistical power for detecting scarce species in community samples, which may lead to a high false-negative rate of species detection. Among the 45 morpho-taxa, 34 (7.12% of the individuals) were false negative, and 25 of these had relatively low representation in the whole community (<6 out of 8,276 individuals). All false negative detections have COI or whole genome sequences in GenBank. It is assumed that these scarce taxonomic groups yielded insufficient amounts of template DNA for PCR amplification and thus failed in the following HTS-based detection. Another likely reason for these false-negative detections is low PCR primer compatibility (Kowalczyk et al., 2011, Deagle et al., 2009, Zhou et al., 2013): species that have binding sites with a low affinity to the primers may capture fewer primer molecules during PCR annealing.

Our metabarcoding approach was able to identify 11 of the morpho-taxa (92.9% of the individuals) in the community samples. However, false negative morpho-taxa (24.4% or 11/45 detected morpho-taxa) was considerably higher than has been reported in previous studies directly comparing morphological and metabarcoding identification, e.g., 0–25.0% or 0/8–2/8 species in Blanckenhorn et al. (2016); between 78-83% species detection in Lobo et al. (2017); 80% or 16/20 of the morphologically identified families in Serrana et al. (2018); and 85% of samples identified at the family level in Carew et al. (2018). Our high missing rate could be due to: 1) we did not control (normalize) the relative abundance of taxa before the HTS library preparation; 2) we used 454 pyrosequencing, which generates smaller numbers of reads than more recent HTS platforms such as Illumina sequencer (Luo et al., 2012); or 3) our macroinvertebrate community sample contained large inter-taxonomic variations in abundance and body mass. If we normalized the relative abundances before HTS library preparation by reducing the sample size of abundant morphological taxa (e.g., Harris et al., 2010), we would expect to obtain a higher detection rate of scarce species. However, the additional processing time required for this might reduce the motivation for using HTS (i.e., time-saving), and this approach also cannot estimate the relative abundances of taxa. It is possible that the recent and rapid advancement of HTS may lead to deeper sequencing technology with a larger read capacity may reduce the missing rate. HTS technology, e.g., 454 sequencing, Illumina or SOLiD has higher error frequency than Sanger sequencing with an approximately 0.5% vs. 0.1% error per nucleotide site, respectively (Shendure 2008). The 454 system tends to cause errors of flame shift and gap opening in samples with low GC contents (Blanckenhorn et al., 2016), which may cause a biased over inflation of the diversity estimates (e.g., Brown et al., 2014, Rosen et al., 2012).

## 5. Conclusion

Reliable and comprehensive, but cost and time efficient methods are of critical importance for biodiversity assessment and environmental monitoring of freshwater ecosystems. In this study, we assess the feasibility of HTS-driven assessments of species diversity and relative abundances. We found supporting evidence for the positive correlation between HTS depth and abundance in the macroinvertebrate community, which had a wider taxonomical range and body mass variation. This finding indicates a possible application of metabarcoding in the quantitative assessment of relative abundances in taxonomically broad communities, although further validations are necessary that use body mass (dry mass) rather than abundance. This correlation also highlighted the pros and cons of the quantitative nature of amplicon-based HTS data. Based on our direct comparison of morphological and metabarcoding analyses, a high rate of false-negative detection was found specifically from scarce species in the community sample. Higher alpha diversity was observed due to the finer taxonomic resolution provided by metabarcoding, and that both incidence and abundance-based estimation of beta diversity reflects that of the morphologically-identified data set. Assessment of the relationship between stream physical-chemical characteristics and the macroinvertebrate taxa detected via morphological and metabarcoding identification showed that both models were significant, but the latter explained a higher percentage of variation which indicates that the metabarcoding approach is capable, and a bit more sensitive in detecting environmental patterns comparable to morphology-based data sets mainly due to its ability to assign finer taxonomic identification (i.e., species level). Technological advancements in both the quantity (i.e., large capacity of total reads) and quality (i.e., low error rate) of HTS would make HTS-based biomonitoring more accurate and reliable for both the research community and the public.

## Acknowledgments

The authors thank Noriyuki Egawa and Yusuke Minakami for their assistance on the fieldwork and laboratory analysis. This research was supported by the Japan Society for the Promotion of Science (JSPS) (grant numbers: 16H04437, 17H01666, 17F17908). M. Gamboa was supported by JSPS Postdoctoral Fellowships for Research in Japan.

## Author Contributions

J. Serrana analyzed and interpreted the data, and wrote the manuscript. Y. Miyake conducted field survey, sample collection, and morphological identification. M. Gamboa conducted molecular laboratory work. K. Watanabe designed the whole study, and interpreted the data. All authors did critical revision and approved the final version of the manuscript.

## References

Andrews S. 2010. FastQC: a quality control tool for high throughput sequence data. Available from: http://www.bioinformatics.babraham.ac.uk/projects/fastqc.

Apha, A., 2005. Wef, 2005 Standard methods for the examination of water and wastewater, 21, pp.258–259.

Aylagas, E., Borja, Á. and Rodríguez-Ezpeleta, N., 2014. Environmental status assessment using DNA metabarcoding: towards a genetics based marine biotic index (gAMBI). PLoS One, 9(3), p.e90529.

Aylagas, E., Borja, Á., Irigoien, X. and Rodríguez-Ezpeleta, N., 2016. Benchmarking DNA metabarcoding for biodiversity-based monitoring and assessment. Frontiers in Marine Science, 3, p.96.

Baird, D.J. and Hajibabaei, M., 2012. Biomonitoring 2.0: a new paradigm in ecosystem assessment made possible by next-generation DNA sequencing. Molecular ecology, 21(8), pp.2039–2044.

Barnes, M.A., and Turner, C.R., 2016. The ecology of environmental DNA and implications for conservation genetics. Conservation Genetics, 17(1), pp.1–17.

Beng, K.C., Tomlinson, K.W., Shen, X.H., Surget-Groba, Y., Hughes, A.C., Corlett, R.T. and Slik, J.F., 2016. The utility of DNA metabarcoding for studying the response of arthropod diversity and composition to land-use change in the tropics. Scientific Reports, 6, p.24965.

Benson, D.A., Cavanaugh, M., Clark, K., Karsch-Mizrachi, I., Lipman, D.J., Ostell, J. and Sayers, E.W., 2012. GenBank. Nucleic acids research, 41(D1), pp. D36–D42.

Blanckenhorn W. U., P. T. Rohner, M. V. Bernasconi, J. Haugstetter, and A. Buser. 2016. Is qualitative and quantitative metabarcoding of dung fauna biodiversity feasible? Environmental Toxicology and Chemistry. 35:1970–1977.

Bucklin, A., Lindeque, P.K., Rodriguez-Ezpeleta, N., Albaina, A. and Lehtiniemi, M., 2016. Metabarcoding of marine zooplankton: prospects, progress, and pitfalls. Journal of Plankton Research, 38(3), pp.393–400.

Carew, M.E., V.J. Pettigrove, L. Metzeling, and A.A. Hoffmann., 2013. Environmental monitoring using next generation sequencing: rapid identification of macroinvertebrate bioindicator species. Frontiers in Zoology, 10: 45.

Carew, M.E., Kellar, C.R., Pettigrove, V.J. and Hoffmann, A.A., 2018. Can high-throughput sequencing detect macroinvertebrate diversity for routine monitoring of an urban river? Ecological indicators, 85, pp.440–450.

Caporaso, J.G., Kuczynski, J., Stombaugh, J., Bittinger, K., Bushman, F.D., Costello, E.K., Fierer, N., Pena, A.G., Goodrich, J.K., Gordon, J.I. and Huttley, G.A., 2010. QIIME allows analysis of high-throughput community sequencing data. Nature methods, 7(5), p.335.

Carrizo, S.F., Jähnig, S.C., Bremerich, V., Freyhof, J., Harrison, I., He, F., Langhans, S.D., Tockner, K., Zarfl, C. and Darwall, W., 2017. Freshwater megafauna: Flagships for freshwater biodiversity under threat. Bioscience, 67(10), pp.919–927.

Clarke, L. J., J. M. Beard, K. M. Swadling, and B. E. Deagle. 2017. Effect of marker choice and thermal cycling protocol on zooplankton DNA metabarcoding studies. Ecology and Evolution, 7: 873–883.

Cowart, D.A., Pinheiro, M., Mouchel, O., Maguer, M., Grall, J., Miné, J. and Arnaud-Haond, S., 2015. Metabarcoding is powerful yet still blind: a comparative analysis of morphological and molecular surveys of seagrass communities. PLoS One, 10(2), p.e0117562.

Deagle B. E., R. Kirkwood, and S. N. Jarman. 2009. Analysis of Australian fur seal diet by pyrosequencing prey DNA in faeces. Molecular Ecology. 18:2022–2038.

Dowle, E. J., X. Pochon, J. C. Banks, K. Shearer, S. A. Wood. 2016. Targeted gene enrichment and high-throughput sequencing for environmental biomonitoring: a case study using freshwater macroinvertebrates, Molecular Ecology Resources, 16: 1240–1254.

Edgar R. C. 2010. Search and clustering orders of magnitude faster than BLAST. Bioinformatics. 26:2460–2461.

Edgar, R. C. 2013. UPARSE: highly accurate OTU sequences from microbial amplicon reads. Nature methods, 10(10), 996.

Elbrecht, V. and F. Leese. 2015. Can DNA-based ecosystem assessments quantify species abundance? Testing primer bias and biomass – sequence relationships with an innovative metabarcoding protocol. PLoSONE, 10: e0130324.

Elbrecht, V., Peinert, B., and Leese, F. 2017. Sorting things out: assessing effects of unequal specimen biomass on DNA metabarcoding. Ecology and evolution, 7(17), 6918–6926.

Elbrecht, V., Vamos, E.E., Meissner, K., Aroviita, J. and Leese, F., 2017. Assessing strengths and weaknesses of DNA metabarcoding-based macroinvertebrate identification for routine stream monitoring. Methods in Ecology and Evolution, 8(10), pp.1265–1275.

Emilson, C.E., Thompson, D.G., Venier, L.A., Porter, T.M., Swystun, T., Chartrand, D., Capell, S. and Hajibabaei, M., 2017. DNA metabarcoding and morphological macroinvertebrate metrics reveal the same changes in boreal watersheds across an environmental variable. Scientific reports, 7(1), p.12777.

Faraway, J.J., 2016. Extending the linear model with R: generalized linear, mixed effects and nonparametric regression models (Vol. 124). CRC press.

Folmer O., M. Black, W. Hoeh, R. Lutz, and R. Vrijenhoek. 1994. DNA primers for amplification of mitochondrial cytochrome c oxidase subunit I from diverse metazoan invertebrates. Molecular Marine Biology and Biotechnology. 3:294–297.

Gibson, J.F., Shokralla, S., Curry, C., Baird, D.J., Monk, W.A., King, I. and Hajibabaei, M., 2015. Large-scale biomonitoring of remote and threatened ecosystems via high-throughput sequencing. PloS one, 10(10), p.e0138432.

González-Tortuero E., J. Rusek, A. Petrusek, S. Gießler, D. Lyras, S. Grath, F. Castro-Monzón, and J. Wolinska. 2015. The quantification of representative sequences pipeline for amplicon sequencing: case study on within-population its1 sequence variation in a microparasite infecting daphnia. Molecular Ecology Resources. 15:1385–1395.

Gotelli, N.J. and Chao, A., 2013. Measuring and estimating species richness, species diversity, and biotic similarity from sampling data.

Hajibabaei M., S. Shokralla, X. Zhou, G. A. C. Singer, and DJ Baird. 2011. Environmental barcoding: A next-generation sequencing approach for biomonitoring applications using river benthos. PLoS ONE. 6:e17497.

Hajibabaei M., J. L. Spall, S. Shokralla, and S. van Konynenburg. 2012. Assessing biodiversity of a freshwater benthic macroinvertebrate community through non-destructive environmental barcoding of DNA from preservative ethanol. BMC Ecology. 12:28.

Harris J. K., J. W. Sahl, T. A. Castoe, B. D. Wagner, D. D. Pollock, and J. R. Spear. 2010. Comparison of normalization methods for construction of large, multiplex amplicon pools for next-generation sequencing. Applied and Environmental Microbiology. 76:3863–3868

Hebert P. D. N., A. Cywinska, S. L. Ball, and J. R. DeWaard. 2003. Biological identifications through DNA barcodes. Proceedings of the Royal Society of London. Series B, Biological sciences. 270:313–321.

Heino, J., 2014. Taxonomic surrogacy, numerical resolution and responses of stream macroinvertebrate communities to ecological variables: are the inferences transferable among regions? Ecological Indicators, 36, 186–194.

Ives, A.R. and Carpenter, S.R., 2007. Stability and diversity of ecosystems. science, 317(5834), pp.58–62.

Jackson, J. K., Battle, J. M., White, B. P., Pilgrim, E. M., Stein, E. D., Miller, P. E., and Sweeney, B. W., 2014. Cryptic biodiversity in streams: a comparison of macroinvertebrate communities based on morphological and DNA barcode identifications. Freshwater Science, 33(1), 312–324.

Kawai T., and K. Tanida. 2005. Aquatic insects of Japan: manuals with keys and illustration (in Japanese). Tokai University Press, Kanagawa, Japan (in Japanese).

Kawanishi R., M. Inoue, M. Takagi, Y. Miyake, and T. Shimizu. 2011. Habitat factors affecting the distribution and abundance of spinous loach, Cobitis shikokuensis, in southwestern Japan. Ichthyological Research. 58:202–208.

Kermarrec L., A. Franc, F. Rimet, P. Chaumeil, J. F. Humbert, and A. Bouchez. 2013. Next-generation sequencing to inventory taxonomic diversity in eukaryotic communities: A test for freshwater diatoms. Molecular Ecology Resources. 13:607–619.

Kowalczyk R., P. Taberlet, E. Coissac, A. Valentini, C. Miquel, T. Kamiński, and J. M. Wójcik. 2011. Influence of management practices on large herbivore diet-case of European bison in Białowieza Primeval Forest (Poland). Forest Ecology and Management. 261:821–828.

Krehenwinkel, H., Wolf, M., Lim, J.Y., Rominger, A.J., Simison, W.B. and Gillespie, R.G., 2017. Estimating and mitigating amplification bias in qualitative and quantitative arthropod metabarcoding. Scientific reports, 7(1), p.17668.

Lobo, J., Shokralla, S., Costa, M.H., Hajibabaei, M. and Costa, F.O., 2017. DNA metabarcoding for high-throughput monitoring of estuarine macrobenthic communities. Scientific reports, 7(1), p.15618.

Lu, H.P., Yeh, Y.C., Sastri, A.R., Shiah, F.K., Gong, G.C. and Hsieh, C.H., 2016. Evaluating community– environment relationships along fine to broad taxonomic resolutions reveals evolutionary forces underlying community assembly. The ISME journal, 10(12), p.2867.

Luo C., D. Tsementzi, N. Kyrpides, T. Read, and K. T. Konstantinidis. 2012. Direct comparisons of Illumina vs. Roche 454 sequencing technologies on the same microbial community DNA sample. PLoS ONE. 7:e30087.

Machado, K.B., Borges, P.P., Carneiro, F.M., de Santana, J.F., Vieira, L.C.G., de Moraes Huszar, V.L. and Nabout, J.C., 2015. Using lower taxonomic resolution and ecological approaches as a surrogate for plankton species. Hydrobiologia, 743(1), pp.255–267.

Macher, J.N., Salis, R.K., Blakemore, K.S., Tollrian, R., Matthaei, C.D. and Leese, F., 2016. Multiple-stressor effects on stream invertebrates: DNA barcoding reveals contrasting responses of cryptic mayfly species. Ecological Indicators, 61, pp.159–169.

Martin, G.K., Adamowicz, S.J. and Cottenie, K., 2016. Taxonomic resolution based on DNA barcoding affects environmental signal in metacommunity structure. Freshwater Science, 35(2), pp.701–711.

Matthaei, C.D., Peacock, K.A. and Townsend, C.R., 1999. Patchy surface stone movement during disturbance in a New Zealand stream and its potential significance for the fauna. Limnology and Oceanography, 44(4), pp.1091–1102.

Menezes, S., Baird, D.J. and Soares, A.M., 2010. Beyond taxonomy: a review of macroinvertebrate trait-based community descriptors as tools for freshwater biomonitoring. Journal of Applied Ecology, 47(4), pp.711–719.

Miyake, Y. and Akiyama, T., 2012. Impacts of water storage dams on substrate characteristics and stream invertebrate assemblages. Journal of Hydro-Environment Research, 6(2), pp.137–144.

Nakagawa H., and Y. Takemon. 2015. Length-mass relationships of macro-invertebrates in a freshwater stream in Japan. Aquat Insects. 36:53–61.

Porazinska D. L., W. Sung, R. M. Giblin-Davis, and W. K. Thomas. 2010. Reproducibility of read numbers in high-throughput sequencing analysis of nematode community composition and structure. Molecular Ecology Resources. 10:666–676.

Ratnasingham S., and P. D. N. Hebert. 2007. BOLD: The Barcode of Life Data System: Barcoding. Molecular Ecology Notes. 7:355–364.

Rosen M. J., B. J. Callahan, D. S. Fisher, and S. P. Holmes. 2012. Denoising PCR-amplified metagenome data. BMC Bioinformatics. 13: 83.

Serrana, J.M., Yaegashi, S., Kondoh, S., Li, B., Robinson, C.T. and Watanabe, K., 2018. Ecological influence of sediment bypass tunnels on macroinvertebrates in dam-fragmented rivers by DNA metabarcoding. Scientific reports, 8(1), p.10185.

Shearer, K. A., Hayes, J. W., Jowett, I. G., and Olsen, D. A. (2015). Habitat suitability curves for benthic macroinvertebrates from a small New Zealand river. New Zealand journal of marine and freshwater research, 49(2), 178–191.

Shendure J., and H. Ji. 2008. Next-generation DNA sequencing. Nature Biotechnology. 26: 1135–1145.

Soininen, E.M., Gauthier, G., Bilodeau, F., Berteaux, D., Gielly, L., Taberlet, P., Gussarova, G., Bellemain, E., Hassel, K., Stenøien, H.K. and Epp, L., 2015. Highly overlapping winter diet in two sympatric lemming species revealed by DNA metabarcoding. Plos One, 10(1), p.e0115335.

Sweeney, B.W., Battle, J.M., Jackson, J.K. and Dapkey, T., 2011. Can DNA barcodes of stream macroinvertebrates improve descriptions of community structure and water quality? Journal of the North American Benthological Society, 30(1), pp.195–216.

Tang C. Q., F. Leasi, U. Obertegger, A. Kieneke, T. G. Barraclough, and D. Fontaneto. 2012. The widely used small subunit 18S rDNA molecule greatly underestimates true diversity in biodiversity surveys of the meiofauna. Proceedings of the National Academy of Sciences. 109:16208–16212.

Thompson, B.W., Riddle, M.J. and Stark, J.S., 2003. Cost-efficient methods for marine pollution monitoring at Casey Station, East Antarctica: the choice of sieve mesh-size and taxonomic resolution. Marine Pollution Bulletin, 46(2), pp.232–243.

Thudi M., Y. Li, S. A. Jackson, G. D. May, and R. K. Varshney. 2012. Current state-of-art of sequencing technologies for plant genomics research. Briefings in Functional Genomics. 11:3–11.

Voelkerding K. V., S. A. Dames, and J. D. Durtschi. 2009. Next-generation sequencing: from basic research to diagnostics. Clinical Chemistry. 55:641–658.

Watanabe K., M. T. Monaghan, Y. Takemon, and T. Omura. 2008. Biodilution of heavy metals in a stream macroinvertebrate food web: evidence from stable isotope analysis. Science of the Total Environment. 394:57–67.

Yu D. W., Y. Ji, B. C. Emerson, X. Wang, C. Ye, C. Yang, and Z. Ding. 2012. Biodiversity soup: metabarcoding of arthropods for rapid biodiversity assessment and biomonitoring. Methods in Ecology and Evolution. 3:613–623.

Zhou X., Y. Li, S. Liu, Q. Yang, X. Su, L. Zhou, M. Tang, R. Fu, J. Li, and Q. Huang. 2013. Ultra-deep sequencing enables high-fidelity recovery of biodiversity for bulk arthropod samples without PCR amplification. GigaScience. 2:4.

Zimmermann, J., Glöckner, G., Jahn, R., Enke, N. and Gemeinholzer, B., 2015. Metabarcoding vs. morphological identification to assess diatom diversity in environmental studies. Molecular Ecology Resources, 15(3), pp.526–542.

